# A versatile ES cell-based melanoma mouse modeling platform

**DOI:** 10.1101/658260

**Authors:** Ilah Bok, Olga Vera, Xiaonan Xu, Neel Jasani, Koji Nakamura, Jordan Reff, Arianna Nenci, Jose G. Gonzalez, Florian A. Karreth

## Abstract

The cumbersome and time-consuming process of generating new mouse strains and multi-allelic experimental animals often hinders the use of genetically engineered mouse models (GEMM) in cancer research. Here, we describe the development and validation of an embryonic stem cell (ESC)-GEMM platform for rapid modeling of melanoma in mice. Our platform incorporates twelve clinically relevant genotypes composed of combinations of four driver alleles (LSL-Braf^V600E^, LSL-Nras^Q61R^, Pten^Flox^, Cdkn2a^Flox^) and regulatory alleles to spatiotemporally control the perturbation of genes-of-interest. Our ESCs produce high contribution chimeras, which recapitulate the melanoma phenotypes of conventionally bred mice. Using our ESC-GEMM platform to modulate Pten expression in melanocytes in vivo, we highlight the utility and advantages of gene depletion by CRISPR-Cas9, RNAi, or conditional knockout for melanoma modeling. Moreover, we use complementary genetic methods to demonstrate the impact of Pten restoration on the prevention and maintenance of Pten-deficient melanomas. Finally, we show that chimera-derived melanoma cell lines retain regulatory allele competency and are a powerful resource to complement ESC-GEMM chimera experiments in vitro and in syngeneic grafts in vivo. Thus, when combined with sophisticated genetic tools, our ESC-GEMM platform enables rapid, high-throughput, and versatile studies aimed at addressing outstanding questions in melanoma biology.

## Introduction

Malignant melanoma is a cancer of melanocytes that is frequently fatal. Despite recent clinical advances in targeted and immune therapy, innate and acquired resistance to treatment necessitates the development of new therapeutic strategies. To this end, novel therapeutic targets must be identified to exploit melanoma-specific genetic dependencies and vulnerabilities. Genomic, genetic, and transcriptomic analyses have revealed numerous genes with putative functions in melanoma initiation, progression, and drug resistance (Berger et al. 2012; Hodis et al. 2012; Krauthammer et al. 2012; 2015; Hayward et al. 2017). Extensive characterization of such candidate driver genes is required to assess their oncogenic potential, unravel the underlying molecular mechanisms, and examine opportunities for therapeutic targeting. Genetically engineered mouse models (GEMMs) that faithfully recapitulate many aspects of the genetics and histopathology of the human malignancy are critical for the success of such efforts (Dow and Lowe 2012; Frese and Tuveson 2007).

Numerous GEMMs to study melanoma have been developed (Pérez-Guijarro et al. 2017), many of which model genetic changes that are frequently observed in human melanoma. The development of transgenic approaches involving the Cre/loxP system enabled the generation of refined melanoma GEMMs, with spatiotemporal control over target gene expression. For instance, oncogenic mutants of Braf and Nras, the most frequently mutated proto-oncogenes in melanoma (Hayward et al. 2017), have been created where the presence of a loxP-flanked transcriptional stop element (LSL) (Perna et al. 2015; Burd et al. 2014) or a wildtype cDNA minigene (Dankort et al. 2007; Mercer et al. 2005) prevent expression of the mutant isoform from the endogenous locus. Similarly, conditional knock-out alleles of the predominant tumor suppressors in melanoma, Pten and Cdkna2, haven been created (Lesche et al. 2002; Krimpenfort et al. 2001). When exposed to a Tamoxifen-inducible Cre allele driven by the Tyrosinase promoter (Bosenberg et al. 2006), the oncogenic Braf^V600E^ and Nras^Q61R^ alleles are expressed at physiological levels while Pten and Cdkn2a are deleted in melanocytes. Combinations of these alleles result in mouse strains that develop melanoma with varying penetrance and latency (Dankort et al. 2009; Dhomen et al. 2009; Burd et al. 2014; Perna et al. 2015). As these GEMMs incorporate the most frequent alterations in human melanoma, they represent excellent “base” models to investigate the functions of additional candidate melanoma genes. Indeed, they have been used to examine the roles of β-catenin, mTorc1, Dnmt3b, Akt, and Trp53 in melanoma progression (Damsky et al. 2011; 2015; Viros et al. 2014; Marsh Durban et al. 2013).

The functions of many genes with putative roles in melanoma remain unexplored. For instance, functional in vivo screens have identified hundreds of candidate genes that may have oncogenic and tumor suppressive effects in melanoma (Karreth et al. 2011; Perna et al. 2015; Mann et al. 2015). Similarly, CRISPR screens identified candidate genes having putative roles in melanoma drug resistance (Shalem et al. 2014; Konermann et al. 2015) and immunotherapy (Manguso et al. 2017). To validate and characterize hits from these screens as well as other genetic and genomic efforts, multi-allelic mouse strains need to be generated. However, creating new mouse alleles and inter-crossing them to generate experimental animals harboring four or more alleles is expensive, slow, and cumbersome, thus rendering conventional mouse modeling an inefficient method to study gene functions in melanoma in vivo. As an alternative to conventional mouse modeling, embryonic stem cell-genetically engineered mouse models (ESC-GEMMs) have been developed (Heyer et al. 2010). This approach relies on GEMM-derived ESCs that are modified in vitro and then used to generate chimeras by blastocyst injection. ESC-derived chimera tissues harbor the alleles of interest, thus enabling the use of chimeras as experimental mice without the need for further breeding (Fig. 1A). When combined with efficient ESC targeting methods and advanced tools for modulating gene expression (Premsrirut et al. 2011; Dow et al. 2014; 2015a), ESC-GEMMs are a powerful strategy for rapid and versatile disease modeling, as has been demonstrated for breast, lung, and pancreatic cancer (Henneman et al. 2015; Zhou et al. 2010; Huijbers et al. 2014; Saborowski et al. 2014).

**Figure 1:**
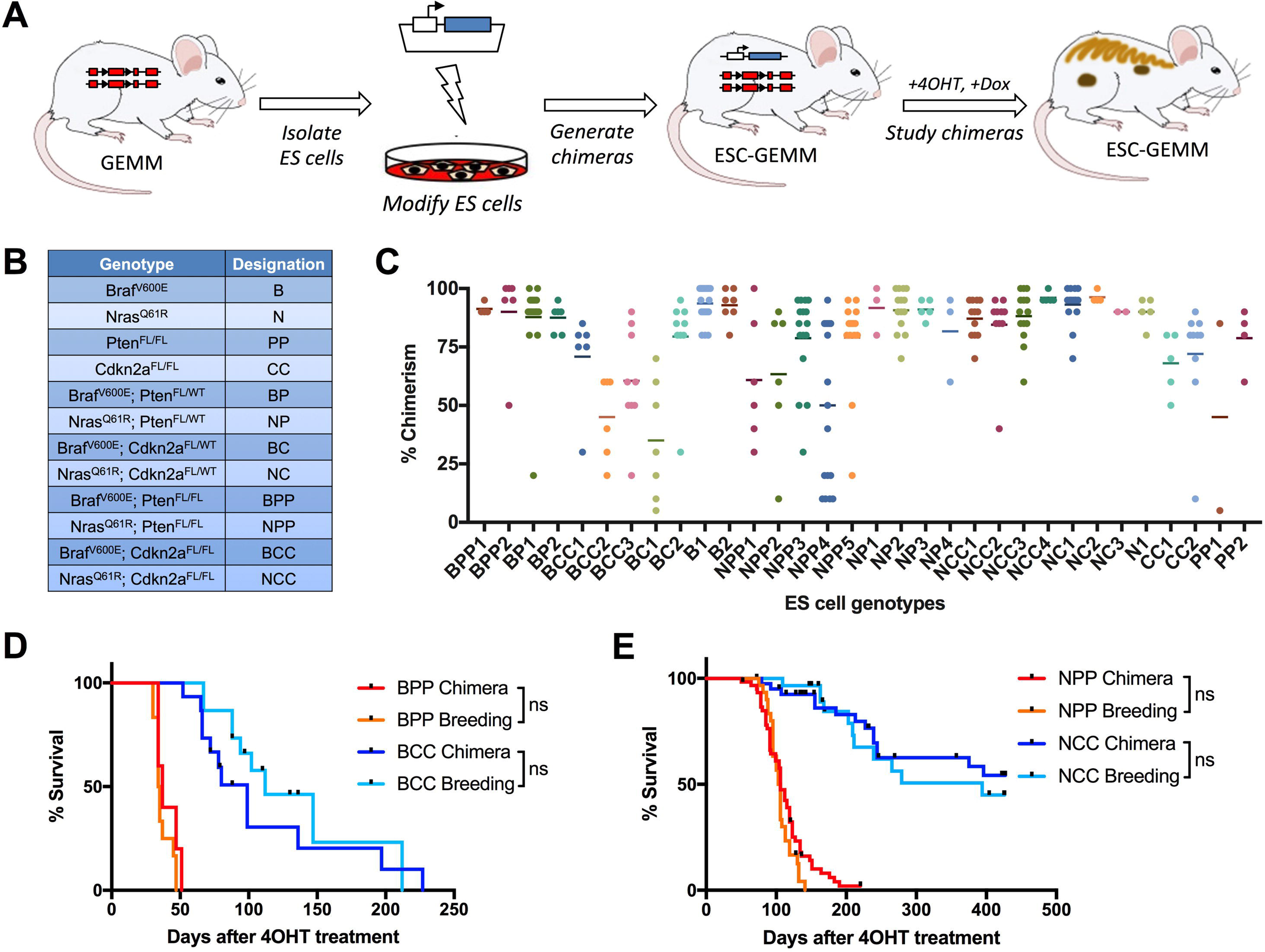
Derivation and validation of ESCs for melanoma mouse modeling. (A) Outline of the ESC-GEMM approach. ESCs are derived from existing GEMMs, which are then modified in vitro. Modified ESCs are used to generate chimeras. ESC-derived melanocytes harbor the alleles required to analyze the phenotype of interest, and chimeras are used as experimental animals. (B) Table showing the different oncogene and tumor suppressor genotype combinations of the newly derived ESC lines. (C) The ability of a panel of ESC lines of the various genotypes to contribute to chimeras. Chimerism is estimated based on the coat color and each dot represent one chimera. (D) Kaplan-Meier comparison of melanoma-mediated reduction in survival of chimeric BPP and BCC (BPP Chimera, n=10; BCC Chimera, n=15) and conventionally bred BPP and BCC mice (BPP Conventional, n=12; BCC Conventional, n=15). (E) Kaplan-Meier comparison of melanoma-mediated reduction in survival of chimeric NPP and NCC (NPP Chimera, n=59; NCC Chimera, n=41) and conventionally bred NPP and NCC mice (NPP Conventional, n=30; NCC Conventional, n=29). Chimeras develop melanoma with the same kinetics and display the same overall survival as conventionally bred animals. ns, not significant.

We report here the generation and validation of an ESC-GEMM platform for melanoma modeling. We derived embryonic stem cell (ESC) lines harboring twelve different combinations of driver alleles as well as alleles to control the modulation of target gene expression. Our ESCs produce high contribution chimeras that exhibit the same melanoma phenotypes as their conventionally bred counterparts. We tested applications of inducible genetic tools and employed them to assess the effects of Pten restoration in Pten-deficient melanomas. Moreover, we have established melanoma cell lines that can complement ESC-GEMM chimera experiments in vitro and in syngeneic grafts in vivo. Our platform is a powerful resource that will accelerate melanoma studies in vivo.

## Results

### Generation and validation of melanoma ESC-GEMMs

GEMMs are critical tools in melanoma research; however, the cost and time investment of generating and maintaining multi-allelic mouse strains precludes a more efficient use of melanoma GEMMs. To alleviate these shortcomings, we created an ESC-GEMM platform for versatile and rapid in vivo melanoma studies. We first intercrossed mice carrying melanoma driver alleles and regulatory alleles to create twelve multi-allelic strains harboring clinically relevant genotypes (Supplemental Table 1). As part of these crosses, we bred our mice to the C57BL/6 background for several generations. We then derived ESC lines from each strain harboring combinations of Cre-inducible Braf^V600E^ (LSL-Braf^V600E^) (Perna et al. 2015) or Nras^Q61R^ (LSL-Nras^Q61R^) (Burd et al. 2014) alleles and conditional knockouts of Pten (Lesche et al. 2002) or Cdkn2a (Krimpenfort et al. 2001) (Fig. 1B). Additionally, our ESCs harbor a melanocyte-specific, 4-Hydroxytamoxifen (4OHT)-inducible Cre recombinase allele (Tyr-CreERt2) (Bosenberg et al. 2006), a Cre-inducible Tet reverse transactivator (CAGs-LSL-rtTA3) (Dow et al. 2014), and a homing cassette in the col1A1 locus (CHC) (Beard et al. 2006) for efficient genomic integration of expression constructs via recombination-mediated cassette exchange (RMCE). This allele combination enables Cre-inducible recombination of driver alleles as well as Cre- and Dox-inducible regulation of genes of interest in melanocytes (Fig. 1A).

ESCs were derived and expanded in 2i media (Ying et al. 2008), and cryopreserved at an early passage (typically between passages 3-5). ESCs were genotyped (Supplemental Table 2) and their sex determined by copy number qRT-PCR for the Y-linked gene Kdm5d. For all twelve genotypes ESC lines having an undifferentiated morphology were selected to test their ability to produce chimeric mice. ESCs were injected into blastocysts derived from Balb/c mice, and blastocysts were transferred to pseudopregnant CD-1 recipient females. All tested ESC lines contributed to chimeras, and most ESCs produced multiple high-contribution chimeric mice having >75% ESC-contribution (Fig. 1C).

We then tested if the ESC-GEMM approach affects melanomagenesis by comparing the phenotypes of chimeras and conventionally bred mice. 3-4 week old chimeras of the various genotypes (see genotype abbreviations in Fig. 1B and Supplemental Table 1) were shaved and topically treated with 25mg/mL 4OHT with a paintbrush on two consecutive days to induce Cre activity in melanocytes. Similarly, 4OHT was administered to conventionally bred LSL-Braf^V600E^; Pten^Flox/Flox^; Tyr-CreERt2 (BPP breeding), LSL-Braf^V600E^; Cdkn2a^Flox/Flox^; Tyr-CreERt2 (BCC breeding), LSL-Nras^Q61R^; Pten^Flox/Flox^; Tyr-CreERt2 (NPP breeding), and LSL-Nras^Q61R^; Cdkn2a^Flox/Flox^; Tyr-CreERt2 (NCC breeding) mice. Importantly, BPP, BCC, NPP, and NCC chimeras developed superficially growing tumors and displayed overall survival rates similar to their conventionally bred counterparts (Fig. 1D,E). Tumors from all genotypes were positive for melanoma markers (Supplemental Fig. S1A), indicating that these were indeed melanomas. Importantly, we observed no tumors in undesired tissues upon gross examination at necropsy. The various genotypes of our ESC-GEMM chimeras led to melanoma phenotypes with different penetrance and latency (Supplemental Fig. S1B,C), and tumor number (Supplemental Fig. S1D). All BPP chimeras and a considerable number of NPP chimeras developed too many tumors for individual tumor nodules to be discerned (denoted as >25 tumors in Supplemental Fig. S1D). When BPP chimeras were treated once with a 10-fold lower dose of 4OHT (2.5mg/mL), melanomas formed with the same latency but at a number low enough to distinguish individual tumors (Supplemental Fig. S1E). The wide-ranging melanoma aggressiveness observed on the various ESC-GEMM backgrounds allows one to choose the model developing the most appropriate phenotype for the question being investigated.

### Testing gene depletion efficiencies in melanoma ESC-GEMMs

Over the last decade our genetic toolbox has significantly expanded, and inducible RNAi and CRISPR technologies have been developed for in vivo use (Premsrirut et al. 2011; Dow et al. 2012; 2015a). We wanted to explore how these technologies can be combined with the ESC-GEMM approach and what their advantages and shortcomings are. We first used the ESC-GEMM approach to assess if gene depletion by the Cre/loxP technology, CRISPR-Cas9, or RNAi provokes similar melanoma phenotypes. We chose to target Pten on a Braf^V600E^ background as this combination results in an aggressive melanoma phenotype (Dankort et al. 2009). We targeted B ESCs with either a Cre- and Dox-inducible Cas9 construct (Han et al. 2017) that also contains a validated sgRNA against Pten (Pten target sequence 1) (Xue et al. 2014) (Fig. 2A, Supplemental Fig. S2A,B). Alternatively, the same B ESCs were targeted with a Dox-inducible, miR30-based RNAi construct containing a validated shRNA targeting Pten (shPten.1523) (Fellmann et al. 2011) (Fig. 2A, Fig. 5A). We also targeted B ESCs with a control sgRNA targeting a non-genic region in chromosome 8 (CR8) (Dow et al. 2015a) or a control shRNA against Renilla Luciferase (shRen.713) (Zuber et al. 2011). For each targeting, several ESC colonies emerged following hygromycin selection and we picked and expanded six to twelve clones each. We confirmed integration of the targeting constructs into the CHC by PCR, which notably was successful in all tested clones. The LSL-Braf^V600E^ allele contains two FRT sites that were included to confirm targeting of the LSL and the V600E mutation to the same chromosome (which was performed in sequential steps) and enable conditional deletion of the mutant allele (Perna et al. 2015). Since the CHC is targeted via FLPe recombinase-dependent RMCE, we had to assess if FLPe-mediated recombination of the LSL-Braf^V600E^ allele had also occurred in targeted ESC clones. We screened targeted clones for LSL-Braf^V600E^ allele recombination by PCR, and excluded all clones in which this recombination event was observed (10-50% of clones).

**Figure 2:**
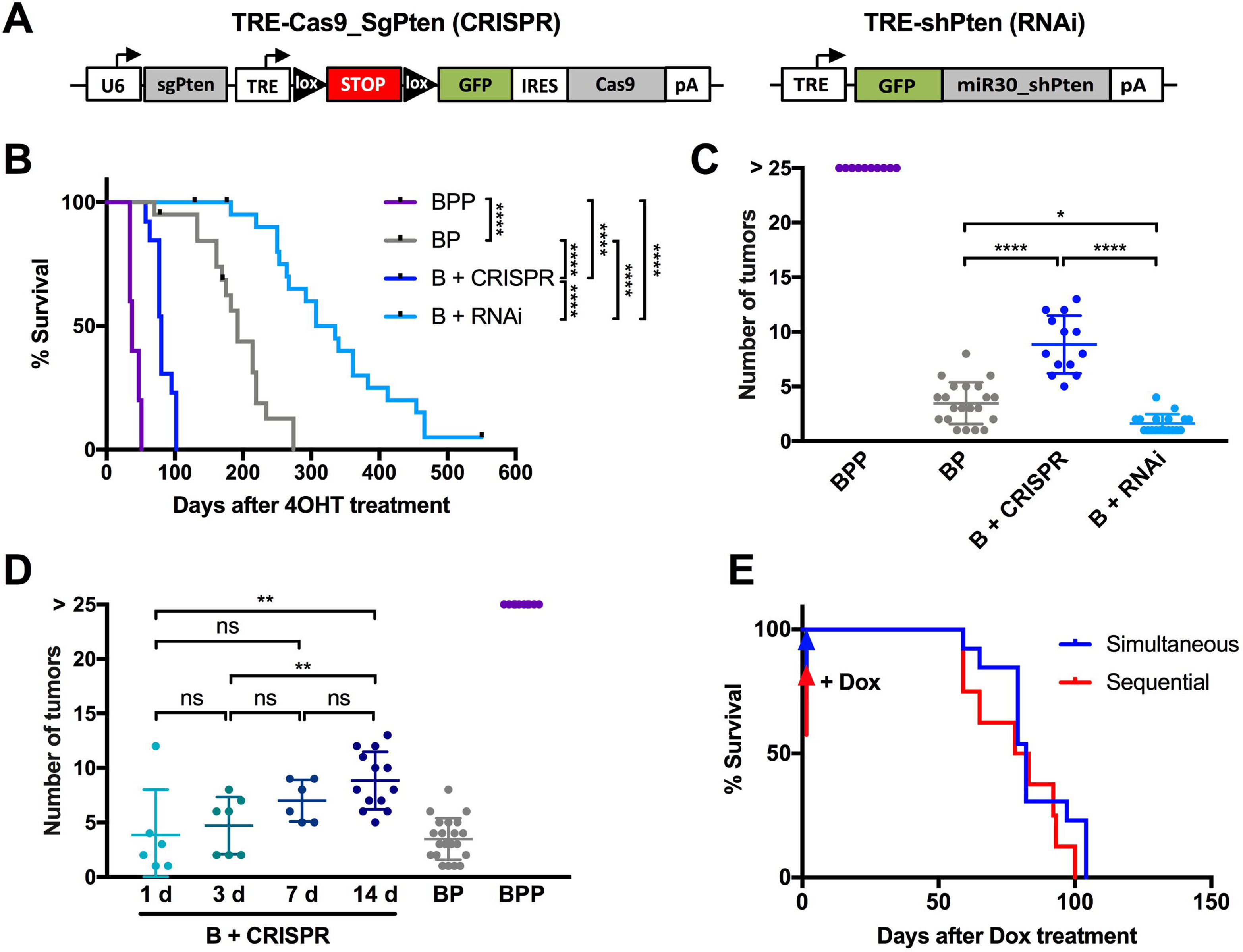
Comparison of the melanoma phenotypes elicited by CRISPR-, RNAi-, and Cre/loxP-mediated gene modulation. (A) Schemes of the CRISPR and shRNA alleles used to delete and silence Pten, respectively. Cas9 is Cre- and Dox-inducible, while shPten is only Dox-inducible. (B) Kaplan-Meier curves comparing the survival of LSL-Braf^V600E^; Pten^FL/FL^ (BPP, n=10) and LSL-Braf^V600E^; Pten^FL/+^ (BP, n=20) chimeras to the survival of B^TRE-Cas9_sgPten^ (B + CRISPR, n=13) and B^TRE-shPten^ chimeras (B + RNAi, n=22) in which Pten was deleted by CRISPR-Cas9 or silenced by RNAi, respectively. (C) Number of melanomas that developed in the chimeras shown in (B). (D) Regulation of melanoma multiplicity with inducible CRISPR-Cas9. Cas9 expression is induced with Dox for 1, 3, or 7 days. The graph shows the number of melanomas that developed in B^TRE-Cas9_sgPten^ chimeras upon the different durations of Cas9 expression. BP, BPP, and B^TRE-Cas9_sgPten^ chimeras (14 days on Dox) from (C) are included for comparison. (E) Temporal control over tumor suppressor deletion with inducible CRISPR-Cas9. Kaplan-Meier analysis shows the survival of B^TRE-Cas9_sgPten^ chimeras in which Cas9 expression was induced for 14 days concomitantly with (Simultaneous, n=13; same cohort as in (B)) or 6 weeks following (Sequential, n=8) 4OHT application. ns, not significant; * p < 0.05; ** p < 0.01; **** p < 0.0001.

Similar to untargeted ESCs, targeted ESCs were used to generate chimeras by injection into Balb/c blastocysts. Importantly, targeted ESCs maintained the ability to produce high-contribution chimeras (Supplemental Fig. 3). We treated 3-4 week old B^TRE-Cas9_sgPten^ and B^TRE-shPten^ chimeras with 4OHT on their back skin to activate Cre in melanocytes. The B^TRE-Cas9_sgPten^ chimeras were fed Dox-containing chow for 14 days immediately following the 4OHT treatment to induce Cas9 expression. B^TRE-shPten^ chimeras were kept on a Dox diet (625mg/kg) continuously to maintain expression of the shRNA and silencing of Pten. Control chimeras (B^TRE-Cas9_sgCR8^ and B^TRE-shRen.713^) treated in a similar fashion did not develop any melanomas (data not shown). We compared melanoma development in the B^TRE-Cas9_sgPten^ and B^TRE-shPten^ chimeras to that in BP and BPP chimeras (generated with untargeted BP and BPP ESCs) where Cre/loxP is used to delete one or both copies of Pten, respectively. Melanomas in B^TRE-Cas9_sgPten^ chimeras developed with a slightly longer latency (Fig. 2B) and a markedly lower multiplicity (Fig. 2C) than in BPP chimeras, suggesting that homozygous deletion of Pten with CRISPR-Cas9 is less efficient than with the Cre/loxP technology. Surprisingly, B^TRE-shPten^ chimeras displayed increased melanoma latency and decreased tumor number compared to all other cohorts (Fig. 2B,C), which was accompanied by an extensive expansion of the cutaneous melanocyte population (Supplemental Fig. S4). Melanomas from B^TRE-Cas9_sgPten^, B^TRE-shPten^, BPP, and BP chimeras displayed reduced expression of Pten and increased phosphorylation of Akt (Supplemental Fig. S5), demonstrating that while all three genetic methods of Pten depletion promote melanomagenesis, their efficiency and phenotype vary drastically.

**Figure 3:**
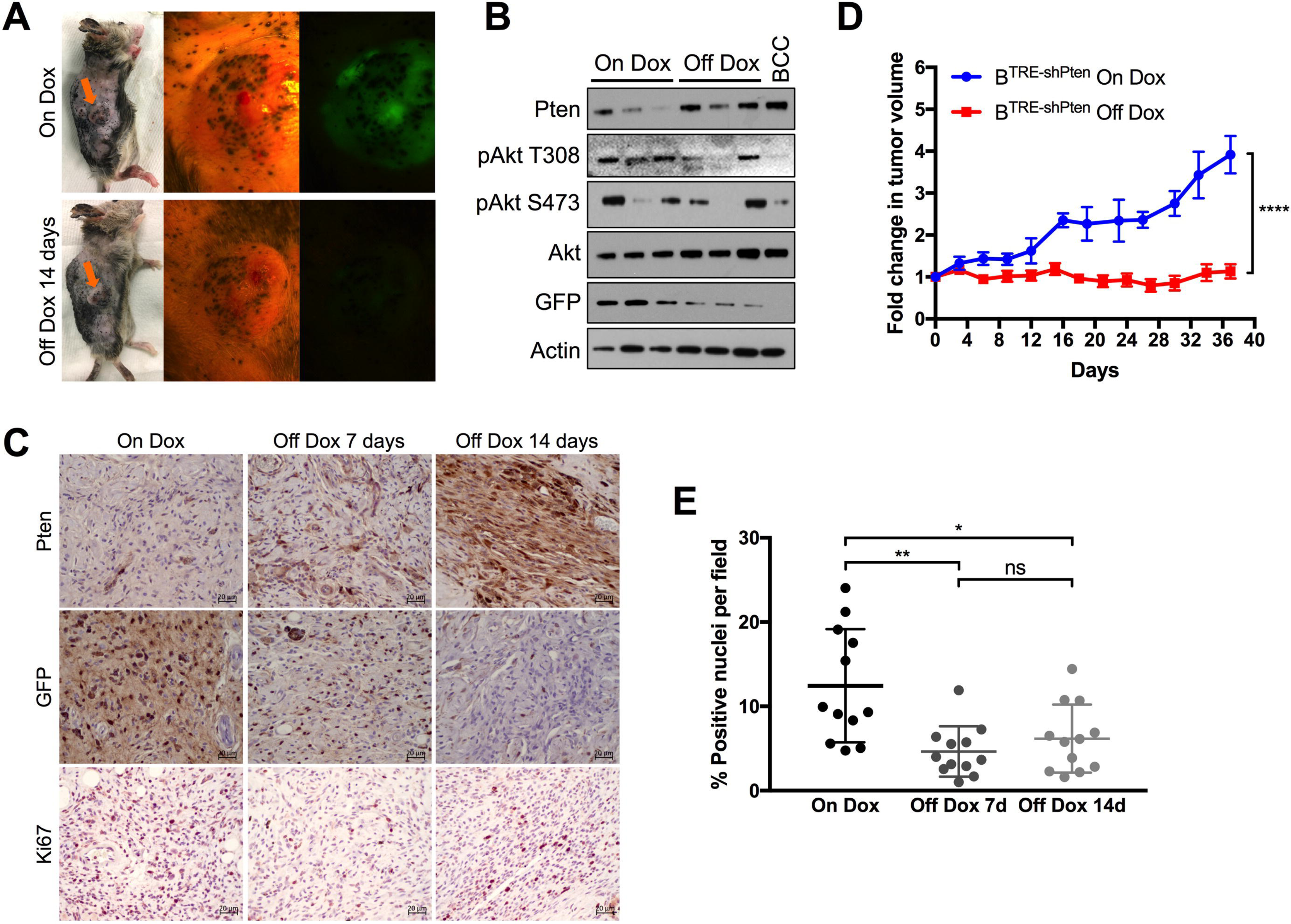
Restoration of endogenous Pten expression in B^TRE-shPten^ chimeras halts melanoma growth. (A) Fluorescent imaging of gross melanomas in B^TRE-shPten^ chimeras where the inducible expression of shPten is linked to GFP. Taking chimeras off Dox for 14 days results in cessation of GFP expression. (B) Western blot showing expression of GFP, Pten, and phosphorylated (pS473 and pT308) and total Akt in melanomas from chimeras on Dox, or taken off Dox for 7 days. Actin was used as a loading control, and a tumor from a BCC chimera is shown for comparison. (C) Immunohistochemistry staining of Pten, GFP and Ki67 on tumors from B^TRE-shPten^ chimeras on Dox or off Dox for 7 or 14 days. (D) Graph displaying the fold change in tumor volume over time in chimeras on Dox compared to chimeras that were taken off Dox on day 0. (E) Quantification of Ki67-positive nuclei per field in tumors from B^TRE-shPten^ chimeras on Dox or off Dox for 7 or 14 days. * p < 0.05; ** p < 0.01; **** p < 0.0001.

### CRISPR applications in melanoma ESC-GEMMs

To further test possible applications for CRISPR-Cas9 in ESC-GEMMs, we took advantage of the Dox-inducibility of our Cas9 construct (Fig. 2A). We first asked whether in addition to varying the 4OHT concentration (Supplemental Fig. S1E) adjusting the duration of Cas9 expression can be used to regulate melanoma multiplicity. We fed B^TRE-Cas9_sgPten^ chimeras a Dox diet for 1, 3, or 7 days and monitored the mice for melanoma development. Indeed, similar to results published in a colon cancer model (Dow et al. 2015a), limiting the duration of Cas9 expression correlated with the formation of fewer melanomas (Fig. 2D, Supplemental Fig. S6A). Notably, the reduction in disease burden had no significant effect on survival (Supplemental Fig. S6B). Thus, by optimizing the duration of Cas9 expression one can adjust the number of tumors without impacting the timing of tumor emergence and the rate of growth.

In the vast majority of multi-allelic GEMMs, all alleles are induced simultaneously. 4OHT treatment of conventional LSL-Braf^V600E^; Pten^FL/FL^; Tyr-CreERt2 mice, for instance, induces recombination of the Braf and Pten alleles resulting in simultaneous expression of Braf^V600E^ and depletion of Pten, respectively. Not having the ability to recapitulate the sequence of mutational events in humans is one of the shortcomings of traditional GEMMs. We tested whether combining our ESC-GEMMs with inducible CRISPR-Cas9 enables sequential modeling of Braf^V600E^ expression and Pten depletion. To this end, we treated B^TRE-Cas9_sgPten^ chimeras with 4OHT after weaning to induce Braf^V600E^ expression and activated Cas9 expression with Dox 6 weeks after 4OHT administration. Notably, delayed Pten deletion provoked a phenotype similar to simultaneous Pten deletion. Melanomas emerged at the same rate, resulting in a similar relative reduction of survival (Fig. 2E, Supplemental Fig. S6C). The number of melanomas was also comparable, although three mice with delayed Pten deletion developed >25 tumors (Supplemental Fig. S6D), possibly due to a Braf^V600E^-mediated expansion of the melanocyte pool in which Pten was deleted. These results demonstrate that inducible CRISPR-Cas9 can be utilized to model the sequential genetic events occurring in human melanomas.

Finally, conventionally bred LSL-Braf^V600E^; Pten^FL/FL^; Tyr-CreERt2 mice often develop spontaneous nevi and melanomas due to leakiness of the Tyr-CreERt2 allele (Damsky et al. 2011). Undesired melanomas may interfere with the experimental read-out and therefore a greater number of mice are needed under certain circumstances. Importantly, spontaneous melanomas did not occur in uninduced B^TRE-Cas9_sgPten^ chimeras (Supplementary Fig. 7), indicating that Cre- and Dox-inducible CRISPR-Cas9 offers tighter spatiotemporal control over tumor suppressor deletion and melanoma induction.

### RNAi-mediated gene silencing in melanoma ESC-GEMMs

The shRNA targeting Pten (shPten.1523) has previously been shown to potently silence Pten expression (Fellmann et al. 2011), and we confirmed highly efficient Pten silencing in vitro and in vivo (Fig. 5A, Supplemental Fig. S5; Supplemental Fig. S13A). Given the potency of shPten.1523, it was surprising that B^TRE-shPten^ chimeras survived significantly longer than BP chimeras in which only one allele of Pten is inactivated (Fig. 2B). Effects of Dox on cancer cells have been widely reported including in melanoma cells (Sun et al. 2009), and we thus tested whether adverse effects of Dox treatment rather than insufficient silencing contributed to the unexpectedly mild melanoma phenotype of B^TRE-shPten^ chimeras. We applied 1μL aliquots of 4OHT to four spots of the back skin of conventionally bred LSL-Braf^V600E^;

Pten^FL/FL^; Tyr-CreERt2 mice to induce focal melanoma development. Immediately following 4OHT treatment, these mice were placed on a 625mg/kg Dox diet or kept on regular chow. Another cohort of mice was given a 200mg/kg Dox diet to explore the effects of a lower Dox concentration. Interestingly, melanomas emerged from almost 90% of 4OHT-treated spots on control and 200mg/kg Dox-treated mice, while only about half of the 4OHT-treated spots in mice given 625mg/kg Dox developed into tumors (Supplemental Fig. S8A). Moreover, Dox had a dose-dependent effect on melanoma size (Supplemental Fig. S8B), demonstrating that melanoma initiation and growth are impaired by Dox. This result may explain, at least in part, the phenotype observed in B^TRE-shPten^ chimeras.

The advantage of Dox-inducible RNAi is that gene silencing is reversible in vivo (Premsrirut et al. 2011), enabling studies into the effect of tumor suppressor restoration (Dow et al. 2015b). To test the reversibility of the inducible shRNA allele and the effect of Pten restoration, we took melanoma-bearing B^TRE-shPten^ chimeras off Dox. Since the expression of shPten.1523 is linked to GFP in our construct, we could observe that GFP-positive tumors became GFP-negative over the course of 14 days of Dox withdrawal (Fig. 3A). Western blots of melanomas on Dox or off Dox for 7 days confirmed a decrease in GFP expression and showed re-expression of Pten accompanied by a decrease in pAkt (Fig. 3B). Similarly, Pten re-expression and a reduction in GFP expression upon Dox withdrawal were observed by immunohistochemistry (Fig. 3C). We also measured melanoma growth in B^TRE-shPten^ chimeras that were either kept on Dox or taken off Dox and, strikingly, observed that Pten restoration completely prevented tumor growth (Fig. 3D, Supplemental Fig. S9A,B). The stalled tumor growth can be attributed to a decrease in proliferation, as shown by fewer Ki67-positive tumor cells, while Dox withdrawal had no effect on the number of TUNEL-positive apoptotic cells (Fig. 3C,E and data not shown). Thus, reversible gene silencing in melanoma ESC-GEMMs demonstrates the requirement for Pten loss for continued tumor growth.

### Inducible expression of cDNAs in melanoma ESC-GEMMs

We decided to further assess Pten re-expression in melanoma by a complementary approach using cDNA expression constructs in our melanoma ESC-GEMMs. To this end, we targeted BPP ESCs with Dox-inducible cDNA constructs encoding Pten or GFP (TRE-Pten and TRE-GFP). We created chimeric mice as described above and applied 4OHT to their back skin to induce Braf^V600E^ expression and delete the floxed Pten alleles. We then placed the BPP^TRE-Pten^ and BPP^TRE-GFP^ chimeras on 200mg/kg Dox food immediately following 4OHT administration to assess if ectopic expression of Pten, but not GFP, can prevent melanoma development. Curiously, the onset of melanoma formation was similar between BPP^TRE-Pten^ and BPP^TRE-GFP^ chimeras (Fig. 4A). However, expression of Pten in BPP^TRE-Pten^ chimeras resulted in increased overall survival (Fig. 4B) and the formation of fewer melanomas than in BPP^TRE-GFP^ control chimeras (Fig. 4C). Immunohistochemistry confirmed the expression of Pten and GFP in melanomas from BPP^TRE-Pten^ and BPP^TRE-GFP^ chimeras, respectively (Fig. 4D). However, only about half of the analyzed melanomas from Dox-treated BPP^TRE-Pten^ chimeras showed strong expression of Pten while almost all tumors from BPP^TRE-GFP^ chimeras exhibited strong GFP staining (Fig. 4E, Supplemental Fig. S10A,B). Interestingly, melanomas from BPP^TRE-Pten^ mice expressed rtTA3 (Supplemental Fig. 10C), indicating that while there is selection against expression of ectopic Pten, this is not due to failed recombination of the CAGs-LSL-rtTA3 allele. Thus, using a complementary cDNA-based approach to study tumor suppressor restoration in ESC-GEMMs, we show that, as expected, Pten reactivation prevents melanomagenesis.

**Figure 4:**
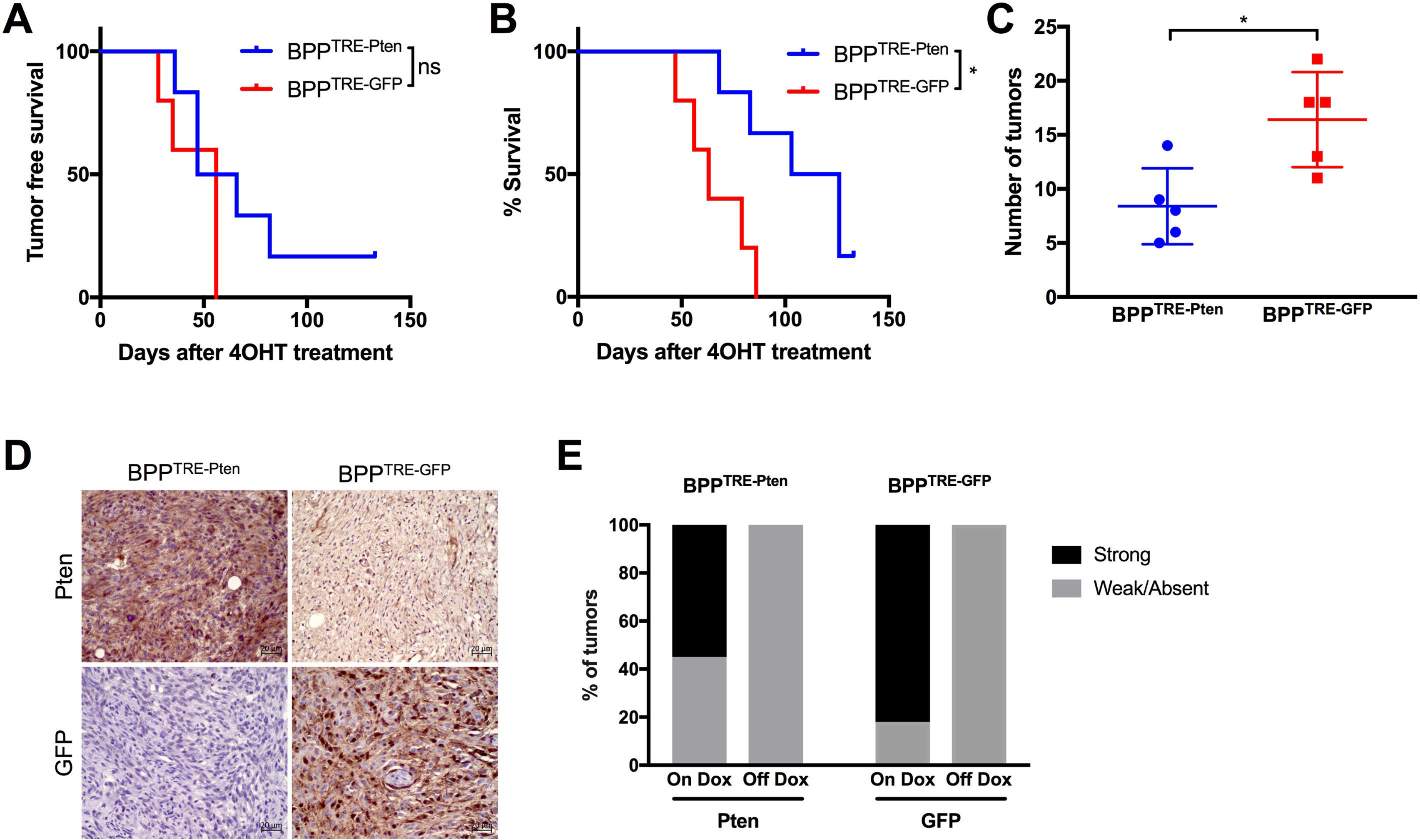
Ectopic Pten re-expression impairs melanoma formation. (A) Kaplan-Meier analysis of tumor-free survival depicting the onset of melanoma formation in BPP^TRE-Pten^ and BPP^TRE-GFP^ chimeras. (B) Kaplan-Meier analysis showing the overall survival of BPP^TRE-Pten^ and BPP^TRE-GFP^ chimeras. (C) Quantification of tumor numbers in BPP^TRE-Pten^ and BPP^TRE-GFP^ chimeras. (D) Immunohistochemistry staining of Pten and GFP in BPP^TRE-Pten^ and BPP^TRE-GFP^ chimeras. (E) Quantification of melanomas with strong or weak/absent Pten staining intensity in melanomas from BPP^TRE-Pten^ and BPP^TRE-GFP^ chimeras. ns, not significant; * p < 0.05.

### ESC-GEMM-derived melanoma cell lines

Murine melanoma cell lines have recently been established from GEMMs (Jenkins et al. 2014; Meeth et al. 2016; Wang et al. 2017). These cell lines can be used for syngeneic transplants in immunocompetent hosts and thus are an invaluable resource to the melanoma and immunotherapy research community. We sought to establish melanoma cell lines from B^TRE-shPten^ and BPP chimeras to further complement our ESC-GEMM approach in general, and the Pten restoration experiments specifically. Intriguingly, our cell lines would have the added advantage of harboring regulatory alleles with which the expression of candidate genes can be modulated. When we prepared melanoma cells from B^TRE-shPten^ and BPP chimeras they ceased to proliferate and became quiescent or senescent within 2-3 passages, similar to previous reports (Jenkins et al. 2014). However, when subcutaneously injected into immunocompromised Nu/Nu mice, tumors readily emerged and we were able to establish cell lines from them (Supplemental Fig. S11A,B). We genotyped all cell lines to confirm they carried the various alleles and had undergone the appropriate Cre-mediated recombinations. Moreover, since our ESCs – and thus melanoma cells derived from them – are on an almost pure C57BL/6 background, we tested if these cell lines would form tumors when syngeneically engrafted. Indeed, tumors readily formed in C57BL/6 recipients (Supplemental Fig. S11B), indicating that these cell lines can be utilized for transplant experiments in the presence of an intact immune system. We then tested if the regulatory alleles present in melanoma cell lines isolated from BPP chimeras (CHC, Tyr-CreERt2, and CAGs-LSL-rtTA3) were still functional and could be used to regulate target gene expression. First, we attempted to target the CHC in melanoma cell lines by RMCE. We used a constitutively active EF1α-GFP expression cassette and were able to generate GFP-positive clones at low efficiency in 6 out of 11 cell lines tested (Supplemental Fig. S12A,B). Correct targeting of the CHC was confirmed by PCR (Supplemental Fig. S12A). Second, we found that none of eleven cell lines tested displayed significant expression of Cre (Supplemental Fig. S12C), and infection with a lentiviral Cre reporter (D’Astolfo et al. 2015) together with 4OHT administration confirmed that Cre had indeed been inactivated (Supplemental Fig. S12D). Given that melanomas in our models are predominantly amelanotic it is likely that the Tyrosinase promoter is inactivated during tumorigenesis. Finally, to validate the functionality of rtTA3, we infected the cell lines with a Dox-inducible GFP lentivirus (TRE-GFP). TRE-GFP cells indeed induced expression of GFP upon Dox administration (Supplemental Fig. S12E,F).

To test the effect of Pten reactivation in B^TRE-shPten^ melanoma cell lines, we first withdrew Dox from the culture media. This restored Pten expression and decreased phosphorylation of Akt (Figure 5A, Supplemental Fig. S13A). As expected, Pten restoration upon Dox withdrawal impaired proliferation and focus formation of B^TRE-shPten^ melanoma cell lines (Fig. 5B,C; Supplemental Fig. S13B,C). We then tested the effect of Pten restoration on the ability of B^TRE-shPten^ melanoma cell lines to form tumors in Nu/Nu mice. We injected cells subcutaneously into three groups of recipients: 1) mice on a Dox diet, 2) mice on a regular diet, and 3) mice on a Dox diet that were switched to a regular diet 18 days following tumor cell transplantation. Tumors in the off Dox cohort (Pten expressed) grew significantly slower than tumors in the on Dox groups (Fig. 5D), resulting in increased survival (Fig. 5E). Interestingly, taking mice bearing established tumors off Dox attenuated tumor growth (Fig. 5D), which also moderately increased survival (Fig. 5E).

**Figure 5:**
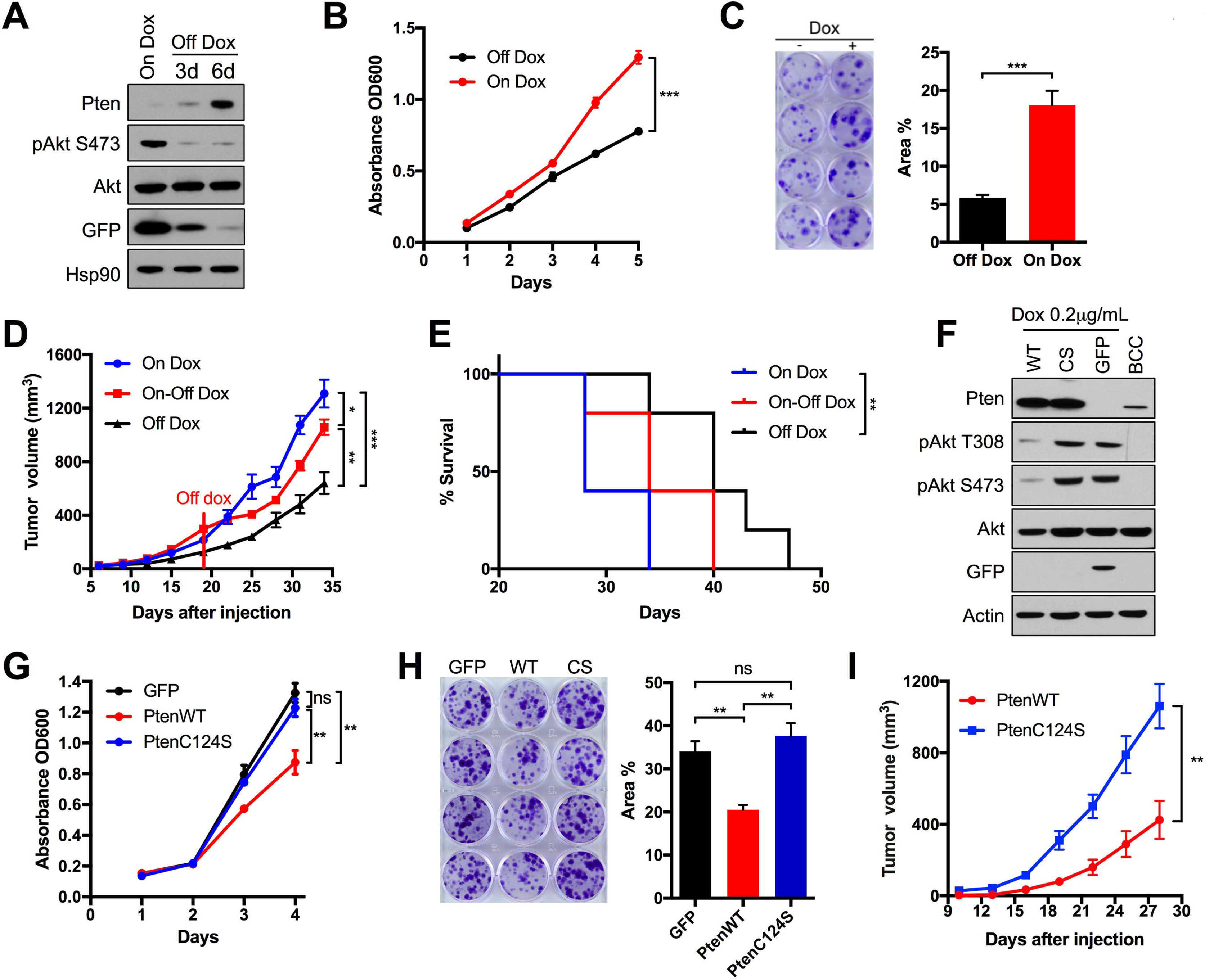
Murine melanoma cell lines as a resource to complement chimera experiments in vitro and in vivo. (A) Melanoma cells from B^TRE-shPten^ chimeras are maintained in Dox-containing media and restore Pten expression upon Dox removal. Western blot shows increased expression of Pten 3 and 6 days after Dox removal, while expression of GFP and AKT pS473 decrease. Total Akt and Hsp90 were used as loading controls. (B) Proliferation curve of B^TRE-shPten^ melanoma cells under on-Dox and off-Dox conditions. (C) Focus formation of B^TRE-shPten^ melanoma cells under on-Dox and off-Dox conditions. (D) Tumor growth of B^TRE-shPten^ melanoma cells in Nu/Nu recipient mice. 5 mice each were kept on a regular diet (Off Dox, n=10 tumors), placed on a Dox diet two days prior to transplantation of tumor cells (On Dox, n=10 tumors), or switched from the On Dox condition to a regular diet 18 days after transplantation of tumor cells (On-Off Dox, n=10 tumors). (E) Survival of Nu/Nu mice shown in (E) bearing B^TRE-shPten^ melanomas (n=5 for each cohort). (F) Western blot of BPP melanoma cells infected with pLenti-TRE-PtenWT, pLenti-TRE-PtenC124S, or pLenti-TRE-GFP lentiviruses. The effect of Dox-mediated induction of Pten expression on Akt phosphorylation (pS473 and pT308) is shown. BCC cells are used as a control for Pten expression, and total Akt and actin were used as loading controls. (G) Proliferation curve of BPP melanoma cells expressing TRE-PtenWT, TRE-PtenC124S, or TRE-GFP (H) Focus formation of BPP melanoma cells expressing TRE-PtenWT, TRE-PtenC124S, or TRE-GFP. (I) Tumor growth of BPP melanoma cells expressing TRE-PtenWT (n=10 tumors) or TRE-PtenC124S (n=10 tumors) transplanted in C57BL/6 mice. Recipient mice were placed on a Dox diet 2 days prior to melanoma cell transplantation. ns, not significant; * p < 0.05; ** p < 0.01; *** p < 0.001.

Next, we re-expressed Pten in BPP melanoma cells using a Dox-inducible lentivirus, pLenti-TRE-PtenWT. As controls, we included a lipid phosphatase-dead Pten mutant (pLenti-TRE-PtenC124S) and GFP (pLenti-TRE-GFP). We observed that stably infected BPP melanoma cells expressed ectopic Pten or GFP upon treatment with Dox (Supplemental Fig. S13G). Only expression of PtenWT, but not PtenC124S or GFP, reduced phosphorylation of Akt (Fig. 5F, Supplemental Fig. S13D), and decreased proliferation and focus formation (Fig. 5G,H; Supplemental Fig. S13E,F). Finally, we transplanted BPP melanoma cells infected with pLenti-TRE-PtenWT or pLenti-TRE-PtenC124S into C57BL/6 recipients. C57BL/6 mice were kept on a Dox diet to induce Pten expression, which led to decreased growth of PtenWT tumors compared to PtenC124S tumors (Fig. 5I). In summary, the melanoma cell lines isolated from chimeras are a useful resource to complement ESC-GEMM experiments and study candidate gene functions in vitro and in vivo.

## Discussion

We have established and validated an ESC-GEMM platform for accelerated melanoma studies in the mouse. Our newly derived ESC lines harbor twelve combinations of alleles modeling the most frequently observed oncogene (BRAF and NRAS) and tumor suppressor (PTEN and CDKN2A) alterations in human melanoma. Chimeras produced with these different ESCs develop melanoma with a wide range of penetrance, latency, and multiplicity. Thus, one can choose the background that is most appropriate for the question being investigated. For instance, the oncogenic potential of candidate oncogenes or tumor suppressors can be readily assessed in low-penetrance/long-latency backgrounds, while putative therapeutic targets can be efficiently evaluated by genetic and pharmacologic means in high-penetrance/short-latency backgrounds. When combined with versatile genetic tools, this platform represents a powerful resource to further unravel the genetic changes underlying melanomagenesis.

In this study, we demonstrate the versatility and flexibility of the melanoma ESC-GEMM platform. Given that it takes less than 2.5 months from ESC targeting to inducing melanomagenesis in experimental chimeras, we anticipate that our platform has the potential to dramatically accelerate melanoma studies in mice. When using the ESC-GEMM approach, several points should be considered. First, while our untargeted ESCs produce high-contribution chimeras, we found that lower-contribution chimeras are more common when targeted ESC clones are used. Importantly, we noted that low chimerism can significantly extend melanoma latency on moderately aggressive backgrounds such as BP (data not shown). Thus, targeted ESCs may produce surplus chimeras that need to be excluded from experiments due to their lower chimerism. As an alternative to injecting ESCs into blastocysts, tetraploid complementation or injecting ESCs into 8-cell embryos could be used. While these techniques produce entirely ESC-derived mice, their success rate is also lower than regular blastocyst injection. Nevertheless, these techniques could be useful when exclusively high-contribution chimeras are needed.

Second, low-contribution chimeras may be useful for certain applications when highly aggressive backgrounds such as BPP are used. We have successfully used chimeras having 5-10% contribution from BPP ESCs, where we induced individual tumors in the ESC-derived skin areas with small aliquots of 4OHT. This approach is practical when individual tumors are to be followed longitudinally. It remains to be determined if such tumors originate from individual transformed melanocytes or if they are polyclonal. In polyclonal tumors, the Tet-ON system may not be operational in all tumor cells as recombination of the CAGs-LSL-rtTA3 allele may fail in tumor subclones. Thus, only a portion of the tumor cells may express or silence the gene of interest. This, in turn, could result in the emergence of “escaper” tumors in prevention or regression studies, where the gene of interest provokes a strong negative selective pressure. For such studies, it is advisable to use high-contribution chimeras and decrease the 4OHT concentration, which considerably reduces the number of melanomas on the BPP background. This increases the likelihood that each tumor originates from individual transformed melanocytes, thereby facilitating the mechanistic analysis of how putative escaper tumors arise. Thus, by varying the 4OHT concentration and means of application, high- and low-contribution chimeras may be used for different experimental contexts.

Third, 2i media containing inhibitors against MEK and GSK3β is ideal for the derivation of ESC lines. However, prolonged culture in 2i media may alter the epigenetic state of ESCs, thereby lowering their developmental potential (Choi et al. 2017). We maintained and targeted our ESCs in 2i media, which may explain why our targeted ESCs produced more low-contribution chimeras. The use of alternative media, such as traditional serum- and LIF-containing media or 2i with alternative inhibitors (Choi et al. 2017), may further improve the potential of our ESCs, but this has to be optimized for each ESC line individually.

We tested whether modulation of gene expression with different technologies – Cre/loxP, CRISPR, and RNAi – produces similar phenotypes. Notably, we found that melanoma latencies and tumor numbers were quite dissimilar. The long latency of melanomas induced by Dox-inducible shRNA was particularly surprising given the potency of shPten.1523 in vitro. While our findings of adverse effects of high Dox concentrations on melanoma initiation and growth provide an explanation for this observation, it remains to be determined if there is a difference in silencing efficiency in tumors compared to cells in culture. Moreover, future studies are required to define the optimal Dox concentration for shRNA experiments where maximum silencing is achieved with minimal effects of Dox, although this will likely be shRNA-dependent. Nevertheless, we highlighted several advantages of Dox-inducible CRISPR-Cas9 and shRNA constructs in combination with the ESC-GEMM approach. While inducible CRISPR-Cas9 can be used to control melanoma multiplicity and the timing of tumor suppressor loss, inducible shRNAs enable reversible gene silencing. We used this approach to restore expression of Pten, which halted melanoma growth. We complemented this approach by inducibly expressing Pten cDNA in BPP chimeras. These models will be useful for evaluating how the effects of Pten restoration compare to inhibition of AKT or mTOR (Dankort et al. 2009).

To complement ESC-GEMM experiments, we isolated murine melanoma cell lines from B^TRE-shPten^ and BPP chimeras. We show that Pten restoration lessens the aggressiveness of these cells in vitro, and diminishes tumor growth in vivo. Murine melanoma cell lines have recently been established and their ability to form tumors when transplanted into immunocompetent syngeneic recipients have made them an invaluable resource to the melanoma community (Jenkins et al. 2014; Meeth et al. 2016; Wang et al. 2017). Similarly, our murine melanoma cell lines form tumors when transplanted into C57BL/6 recipients. Importantly, however, transgenic cassettes can be incorporated into the CHC via RMCE in BPP cells and their expression can be controlled using the integrated Tet-ON system. Thus, cancer gene function can be readily modulated and assessed in vitro and in vivo. Moreover, given the recent success of immunotherapy, we foresee immense utility of these cells for studies focused on delineating the molecular mechanisms governing melanoma immune suppression. In summary, we have generated and validated an ESC-GEMM platform consisting of a panel of newly derived ESCs and murine melanoma cell lines that will considerably facilitate future in vivo melanoma studies.

## Materials and Methods

### ESC line generation

Approximately 28-day-old females were injected with 5U pregnant mare serum gonadotropin (PMSG, Sigma Aldrich, Cat. # G4877-2000IU) and with 5U human chorionic gonadotropin 48 hours later. Hormone-injected females were paired with stud males overnight, and females were euthanized at 3.5 days post coitum. Uterine horns were flushed with EmbryoMax Advanced KSOM Embryo Medium (Millipore, Cat. # MR-101-D) and blastocysts and late morulas were collected in EmbryoMax Advanced KSOM Embryo Medium. Embryos were incubated in humidified incubator (37°C, 5% CO2) for 4-6 hours to allow morulas to develop to blastocysts. Blastocysts were then plated on the gelatinized 4-well plates with mitotically inactivated DR4 mouse embryonic fibroblasts (MEFs) in 2i medium and incubated until ESC colonies formed. ESC colonies were picked with a P20 pipette in 20 μL PBS and dissociated in 20μl 0.05% trypsin (Life Technologies, Cat. # 25-300-120). Single cells were plated on gelatinized 24-well plates with and without DR4 feeders for expansion and for DNA isolation, respectively. ESC clones were genotyped (genotyping primers are listed in Supplemental Table 2) and the sex was determined using RT-PCR to detect Y-linked Kdm5d (Thermo Fisher, Assay ID Mm00528628_cn), Tfrc was used as internal control (Thermo Fisher, Cat. # 4458366).

### ESC targeting and genotyping of targeted ESCs

ESCs were trypsinized, washed in PBS, and .3×10^6^ cells were collected by centrifugation. Supernatant was aspirated and cells were carefully resuspended in P3 solution (Lonza, Cat. # V4XP-3012) containing the 20μg targeting vectors and 10μg pCAGGS-FLPe recombinase expression vector. Cells were transferred to a cuvette and electroporated using the Lonza 4D-Nucleofactor on the “ES, mouse” setting. Nucleofected ESCs were transferred to gelatinized 6 cm dishes with DR4 feeders in 2i media. ESCs were split 24 hours following the nucleofection onto one 10 cm dish. Selection with hygromycin (100-125 mg/mL, VWR, Cat. # 97064-454) was initiated 48 hours following nucleofection. Up to 12 surviving clones were picked after 7-10 days, expanded, and genotyped to validate successful targeting vector integration into the CHC.

### ESC preparation and injection

ESCs were trypsinized and replated on gelatinized 6 well plates and incubated for 30 minutes for feeder depletion. ESCs were gently washed off the plate twice with 1mL of ESC medium and each wash was collected in separate 15mL conical tubes. Cells were collected by centrifugation and resuspended in M2 medium (Sigma-Aldrich, Cat. # M7167) (400μL for the first wash, 200μL for the second wash). The Gene Targeting Core at Moffitt Cancer Center performed injections into blastocyst isolated from Balb/c females, and transferred blastocysts into pseudo-pregnant CD-1 females.

### Plasmids

The empty Dox-inducible cDNA, CRISPR-Cas9 and RNAi vectors for CHC targeting as wells as pCAGGS-FLPe were described previously (Premsrirut et al. 2011; Beard et al. 2006; Dow et al. 2015a) and provided by L. Dow. The targeting vectors used were col1a1-TRE-GFP-miRE_shRen.713, col1a1-TRE-GFP-miRE_shPten.1523, col1a1-TRE-LSL-GFP-IRES-Cas9_U6-sgPten, col1a1-TRE-LSL-GFP-IRES-Cas9_U6-sgCR8, col1a1-TRE-Pten, col1a1-TRE-GFP. The Cre reporter was a gift from N. Geijsen (Addgene plasmid #62732). col1a1-EF1 -GFP was generated by removing TRE from col1a1-TRE and GFP from pLenti-GFP (Addgene plasmid #17448) and the EF1α core promoter from lentiCRISPRv2_puro (from T. Jacks) were inserted using standard cloning techniques. pLenti-TRE-GFP-Blast was generated by replacing the CMV promoter in pLenti-GFP-Blast with the TRE3G promoter from pRRL-TRE3G-GFP-PGK-Puro-IRES-rtTA3 (from J. Zuber). pLenti-TRE-PtenWT-Blast and pLenti-TRE-PtenC124S-Blast were created by replacing GFP in pLenti-TRE-GFP-Blast with mouse Pten wildtype or mutant cDNA.

### Mouse strains

All animal experiments were conducted in accordance with an IACUC protocol approved by the University of South Florida. All mouse strains and alleles used in this study have been described previously: LSL-Braf^V600E^ (Perna et al. 2015), LSL- Nras^Q61R^ (Burd et al. 2014), Pten^Flox^ (Lesche et al. 2002), Cdkn2a^Flox^ (Krimpenfort et al. 2001), Tyr-CreERt2 (Bosenberg et al. 2006), CAGs-LSL-rtTA3 (Dow et al. 2014), CHC (Beard et al. 2006). Chimerism was determined by the percentage of brown/black fur. Chimeras with at least 75% ESC contribution were used for almost all experiments.

### Tumor induction and measurement

Mice were put under anesthesia and back hair was shaved with clippers. 4OHT (Sigma Aldrich, Cat. # H6278-50MG) dissolved in DMSO was administrated with a paintbrush on the back skin of 3-4 week old mice on two consecutive days and was used at 25mg/mL unless noted otherwise. Doxycycline feed (200 mg/kg or 625 mg/kg) was purchased from Envigo and replenished twice weekly. Tumors were measured using calipers and volumes calculated using the formula (width^2^ x length)/2.

### Mouse melanoma cell line generation

Tumors were harvested, washed in 70% ethanol for 10 seconds, and washed in PBS. Tumor tissues were chopped into small pieces, collected in a 50mL conical tube, and washed with 20-25mL RPMI with 5% FBS and 1% penicillin/streptomycin. Tissue pieces were spun down at 1500 rpm for 5 minutes and incubated with 1mg/mL Collagenase/Dispase (Sigma-Aldrich, Cat. # 10269638001) in RPMI with 5% FBS and 1% penicillin/streptomycin for 20 minutes in a humidified incubator at 37 °C. Tissues were washed with PBS and further dissociated in 0.25% Trypsin (VWR, Cat. # VWRL0154-0100) for 30 minutes in the incubator. Tissues were spun down and washed in PBS, and plated in RPMI with 5% FBS and 1% penicillin/streptomycin. Cell line stocks were cryopreserved at passage 2. To establish BPP lines, melanoma cells were passaged through athymic nude mice (The Jackson Laboratory, J:NU 007850) after 2 passages in vitro. Subcutaneous tumors were harvested from nude mice and processed for tissue culture as described above.

### Targeting of mouse melanoma cell line

For M10M7, 0.3×10^6^/plate cells were seeded in three 10 cm dishes. For M10M1 and M10M3, 0.5×10^6^/plate cells were seeded in two 10 cm dishes. Cells were co-transfected with pCAGGS-FLPe and col1a1-EF1α-GFP at a 1:2 ratio using FuGENE HD Transfection Reagent (Promega, Cat. # E2311), and selected in 50μg/mL hygromycin starting 48 hours post-transfection. GFP-positive colonies were picked after three weeks.

### Fluorescent dissecting microscopy

Mice were put under anesthesia and hair removal cream was applied on the back. Hair removal cream was removed after 1-2 minutes and skin carefully rinsed with PBS. Fluorescent images of mouse tumors were taken with a Leica DFC450C microscope.

### RNA isolation, cDNA synthesis, and RT-qPCR

Tumor tissues were harvested and snap frozen in LN_2_. Tumor tissues were homogenized with zirconium beads (Benchmark Scientific, Cat. # D1032-30) and RNA was isolated using the RNeasy Plus Mini Kit (Qiagen, Cat. # 74134) as per the manufacturer’s protocol. cDNA was synthesized using PrimeScript RT Master Mix (Takara Bio, Cat. # RR036A) per the manufacturer’s protocol. qPCRs were performed using primers specific to rtTA3 with PerfeCTa SYBR Green FastMix (QuantaBio, Cat. # 95073-012). Mouse β-actin was used as a loading control.

### T7 endonuclease assay

Genomic DNA was isolated and region of sgPten targeting was PCR amplified using GoTaq G2 Green Master Mix (Promega, Cat. # M7823). The PCR product was purified using E.N.Z.A. Gel Extraction Kit (Omega Bio-Tek, Cat. # D2500-02). 200 ng of purified PCR product were denatured and re-annealed in NE Buffer 2 (New England BioLabs, Cat. # B7002S). T7 endonuclease 1 (New England BioLabs, Cat. # M0302S) was added and incubated at 37°C for 15 minutes. Samples were separated on a 2.5% agarose gel and imaged using a Bio-Rad Gel Doc.

### Proliferation assay and colony formation assay

Cells were cultured in RPMI with 5% FBS and 1% penicillin/streptomycin. For proliferation assay, 1×10^3^ cells were seeded in 96-well plates in triplicates and harvested for five consecutive days. Cells were fixed in cold 4% paraformaldehyde (VWR, Cat. # 101176-014) for 10 minutes, washed twice with PBS, and stained with 0.5% crystal violet (VWR, Cat# 97061-850) solution in 25% methanol for an hour. Stained cells were washed with DI water and dried overnight. Crystal violet was extracted with 10% acetic acid and absorbance at 600 nm was measured using a plate reader. For colony formation assay, 400 cells were seeded in 12-well plates in triplicates and harvested after 10-14 days. Cells were fixed and stained using the same procedure as for proliferation assays, and the percent area of crystal violet staining was quantified using Image J.

### Immunoblotting

Cells were washed with cold PBS and lysed in cold RIPA buffer containing a protease inhibitor cocktail (Sigma-Aldrich, Cat. # 4693124001) and phosphatase inhibitor cocktail (Cell Signaling Technology, Cat. # 5870S). Tumor tissues were harvested and snap frozen in LN_2_ until ready to be lysed. Tissues were dissociated in RIPA buffer with protease inhibitor and phosphatase inhibitor with pellet pestle (Sigma-Aldrich, Cat. # Z359947-100EA) followed by sonication for 15 minutes. All primary antibodies were incubated overnight at **°**C. Antibodies for phosphorylated protein was diluted in Tris-buffered saline (TBS)/0.1% Tween-20/3% BSA, and all other antibodies in TBS/0.1% Tween20/5% milk. Pten (1:2,000, Cell Signaling Technology, Cat. # 9188S), pAkt T308 (1:500, Cell Signaling Technology, Cat. # 13038T), pAkt S473 (1:2,000, Cell Signaling Technology, Cat. # 4060S), AKT (1:5,000, Cell Signaling Technology, Cat. # 4691T), GFP (1:2,000, Cell Signaling Technology, Cat. # 2956S), Hsp90 (1:5,000, Cell Signaling Technology, Cat. # 4874), beta-actin (1:10,000, Fisher Scientific, Cat. # AM4302) were used.

### Immunohistochemistry

Tumor tissues were harvested and fixed in 10% neutral buffered formalin overnight. Tissues were dehydrated in 70% ethanol the following day. IDEXX BioAnalytics (Columbia, MO) performed paraffin embedding, sectioning, hematoxylin and eosin staining, Ki67 immunohistochemistry, and TUNEL staining. Anti-S100 (1:250, Dako, Cat. # Z0311), anti-MART1 (1:250, Sigma-Aldrich, Cat. # SAB4500949-100UG), Pten (1:125, Cell Signaling Technology, Cat. # 9188S), and GFP (1:200, Cell Signaling Technology, Cat. # 2956S) were used for immunohistochemistry.

### Lentiviral transduction

HEK293 Lenti-X (Clontech, Cat. # 632180) cells were cultured in DMEM with 10% FBS. Second generation lentivirus was generated using 0.66 μg VSVG, 5.33 μg Δ8.2, and 6 μg lentiviral construct per 10 cm dish. Transfection was performed using JetPRIME (VWR, Cat. # 89129-924). Lentiviral titers were estimated using Lenti-X GoStix Plus (Takara Bio, Cat. # 631281). Lentiviral transduction was performed overnight in the presence of 8μg/mL polybrene (Sigma-Aldrich, Cat. # 107689-10G).

### Statistical analysis

Statistical analysis was performed using GraphPad Prism software. Survival data were compared by applying the Log-rank (Mantel-Cox) test, and all other data were analyzed with the unpaired two-tailed t-test or ordinary one-way ANOVA. A p-value of 0.05 was considered statistically significant.

## Acknowledgements

We are grateful to G. DeNicola for critical reading of the manuscript. We thank D. Tuveson, N. Sharpless, H. Wu, A. Berns, S. Lowe, M. Bosenberg, and R. Jaenisch for mouse alleles, and L. Dow, J. Zuber, and T. Jacks for plasmids. This work was supported by grants to F.A.K. from the NIH/NCI (K22 CA197058, R03 CA227349), the Melanoma Research Alliance (MRA Young Investigator Award), the American Cancer Society (IRG-14-189-19), the Moffitt Skin SPORE (P50 CA168536) Career Development Program, a Miles for Moffitt Milestone Award, and a Harry J. Lloyd Charitable Trust Career Development grant.

## Author Contributions

IB designed and performed experiments, and analyzed data. OV, XX, KN, NJ performed in vitro experiments. JR performed in vivo experiments. AN derived and targeted ESCs. JGG performed blastocyst injections. FAK supervised the study, designed experiments, and analyzed data. IB and FAK wrote the manuscript with input from all authors.

