## Supplemental Figure Legends for "A versatile ES cell-based melanoma mouse modeling platform"

**Supplemental Figure S1**: Comparison of melanomas that arise in chimeras from the various genotypes. (**A**) The tumors arising in BPP, BP, BCC, BC, NPP, NP, and NCC chimeras stain positive for a marker cocktail consisting of S100 and MART1 by immunohistochemistry, confirming that these tumors are melanomas. (**B**) Kaplan-Meier survival analysis of BPP, BP, BCC, BC, and B chimeras. Note that the survival curves of BPP and BCC chimeras are also shown in Figure 1D. (**C**) Kaplan-Meier survival analysis of NPP, NP, NCC, NC, and N chimeras. Note that the survival curves of NPP and NCC chimeras are also shown in Figure 1E. (**D**) Graph showing the number of melanomas in BPP, BP, BCC, BC, NPP, NP, NCC, NC, B, and N chimeras. Note that NC, B, and N chimeras did not develop melanomas during the course of the study. (**E**) Comparison of the number of melanomas that BPP chimeras develop when treated with 25mg/mL or 2.5mg/mL 4OHT.

**Supplemental Figure S2**: Validation of the Pten-targeting sgRNA. (**A**) A lentivirus (lentiCRISPRv2_puro from T. Jacks) containing Cas9 and a Pten-targeting sgRNA from Xue et al. (Xue et al. 2014) was used to infect the murine melanoma cell line T10921. A sgRNA targeting a non-genic region on chromosome 8 (CR8) was used as a negative control (Dow et al. 2015). Western blot of lentivirus-infected polyclonal T10921 cells showed that sgPten reduced Pten expression. (**B**) T7 Surveyor assay on the polyclonal T10921 cells shown in (A).

**Supplemental Figure S3**: Graph displaying the extent of chimerism of mice generated with targeted ESCs. Each dot represents a chimera.

**Supplemental Figure S4**: Appearance of B^TRE-shPten^ chimeras. (**A**) Examples of B^TRE-shPten^ chimeras that were treated with 4OHT and then fed a Dox diet for several months. Note the marked hyperpigmentation on areas of the back skin where 4OHT was applied. (**B**) H&E stains of skin and melanomas from B^TRE-shPten^ and BPP chimeras showing the extent of melanocyte expansion.

**Supplemental Figure S5**: Pten expression and Akt activation in melanomas driven by CRISPR-, RNAi-, or Cre/loxP-mediated deletion of Pten. (**A**) Western blot showing expression of Pten, phosphorylated (S473, T308) and total Akt, and GFP in melanomas from BP, BPP, B^TRE-Cas9_sgPten^, and B^TRE-shPten^ chimeras. Hsp90 was used as a loading control, and melanomas from BCC chimeras are shown for comparison. (**B**) Pten immunohistochemistry on melanomas from BP, BPP, B^TRE-Cas9_sgPten^, and B^TRE-shPten^ chimeras. A melanoma from a BCC chimera is shown for comparison.

**Supplemental Figure S6**: Applications of inducible CRISPR-Cas9. (**A**) Images of B^TRE-Cas9_sgPten^ chimeras that were fed a Dox-containing diet for 1, 3, or 7 days immediately following 4OHT application. Images were taken 6 weeks post 4OHT application, and arrows indicate nevi (yellow) and emerging melanomas (orange). (**B**) Kaplan-Meier survival curve of B^TRE-Cas9_sgPten^ chimeras fed Dox for 1, 3, 7, or 14 days. A control sgRNA targeting a non-genic region on chromosome 8 (CR8) has no effect on survival. (**C**) Kaplan-Meier survival curve of B^TRE-Cas9_sgPten^ chimeras in which Cas9 expression was activated for 14 days immediately following 4OHT treatment (Simultaneous) or 6 weeks later (Sequential). This graph is the same as in Figure 2E, except that the X-axis displays Days after 4OHT treatment. (**D**) The number of melanomas in the B^TRE-Cas9_sgPten^ chimeras shown in Figure 2E. ns, not significant; * *p* < 0.05; ** *p* < 0.01.

**Supplemental Figure S7**: Comparison of the number of nevi and melanomas that spontaneously develop in conventionally bred LSL-Braf^V600E^; Pten^FL/FL^; Tyr-CreERt2 mice (BPP1-9) or B^TRE-Cas9_sgPten^ chimeras (sgPten1-6) in the absence of 4OHT or Dox administration.

**Supplemental Figure S8**: Effects of Dox on melanoma development. (**A**) Melanoma development was induced on 4 individual spots of the backs of conventionally bred LSL-Braf^V600E^; Pten^FL/FL^; Tyr-CreERt2 mice using 1μL aliquots of 4OHT. Mice were switched to Dox-containing diets (200mg/kg or 625mg/kg) immediately following 4OHT application or kept on a regular diet. The graph shows the percent of 4OHT-induced spots that formed melanomas. (**B**) Tumor size of the LSL-Braf^V600E^; Pten^FL/FL^; Tyr-CreERt2 mice from (A) after 6 weeks on Dox. ns, not significant; * *p* < 0.05; ** *p* < 0.01.

**Supplemental Figure S9**: Pten restoration in B^TRE-shPten^ chimeras. (**A**) Fold change in volume of individual melanomas in B^TRE-shPten^ chimeras that were continuously kept on a Dox diet. (**B**) Fold change in volume of individual melanomas in B^TRE-shPten^ chimeras that were taken off Dox. The average melanoma volumes of each cohort are shown in Figure 3D.

**Supplemental Figure S10**: Pten expression in BPP^TRE-Pten^ chimeras. (**A**) Images of individual tumors from BPP^TRE-Pten^ chimeras (on Dox and off Dox) stained for Pten by immunohistochemistry. (**B**) Images of individual tumors from BPP^TRE-GFP^ chimeras (on Dox and off Dox) stained for GFP by immunohistochemistry. (**C**) Analysis of rtTA3 expression in melanomas from BPP^TRE-Pten^ and BPP^TRE-GFP^ chimeras by real-time PCR.

**Supplemental Figure S11**: Melanoma cell line derivation from ESC-GEMM chimeras. (**A**) Schematic outline of the procedure to establish BPP and B^TRE-shPten^ melanoma cell lines from EC-GEMM chimeras. (**B**) Established melanoma cell lines form tumors when subcutaneously transplanted into immunocompromised Nu/Nu and immunocompetent C57BL/6 recipient mice.

**Supplemental Figure S12**: Testing the functionality of regulatory alleles in ESC-GEMM-derived melanoma cell lines. (**A**) Targeting vectors can be inserted into murine melanoma cell lines by recombination mediated cassette exchange. BPP melanoma cell lines were co-transfected with a targeting vector containing a constitutive EF1α-GFP construct and a pCAGGS-FLPe expression plasmid. A stable clone expressing GFP is shown in the upper panel. The lower panel shows a genotyping PCR of 3 clones confirming that the EF1α-GFP construct was inserted into the CHC locus by RMCE. (**B**) The number of GFP-positive clones that emerged from 900,000 cells following EF1α-GFP and pCAGGS-FLPe transfection and selection in Hygromycin. Each dot represents a transfection experiment. (**C**) Tyr-CreERt2 is silenced in BPP melanoma cell lines. Western blot for Cre shows that Tyr-CreERt2 is not expressed in the murine melanoma cell lines. A nevus isolated from a BPP mouse served as positive control. (**D**) A BPP melanoma cell line was infected with a Cre-dependent RFP/GFP lentiviral reporter. The cells were then either exposed to 4OHT or infected with Adenoviral-Cre (positive control) to induce reporter recombination. 4OHT treatment failed to induce reporter recombination, demonstrating that Tyr-CreERt2 is not active. (**E**) The Tet-On system is functional in murine melanoma cell lines. Western blot showing GFP expression upon Dox treated in BPP melanoma cells infected with a lentivirus containing a TRE-GFP construct. (**F**) Immunofluorescence images of the cells shown in (E).

**Supplemental Figure S13**: The effect of Pten restoration in murine melanoma cell lines. (**A-C**) A second B^TRE-shPten^ cell line was shown to restore Pten expression (A), and reduce proliferation (B) and focus formation (C) upon Dox withdrawal. (**D-F**) A second BPP cell line infected with TRE-PtenWT, TRE-PtenC124S, or TRE-GFP is shown. Induction of PtenWT, but not PtenC124S or TRE-GFP, repressed Akt phosphorylation (D) and reduced proliferation (E) and focus formation (F). (**G**) Western blot for Pten and GFP to show the Dox-inducible nature of the TRE-PtenWT, TRE-PtenC124S, and TRE-GFP lentiviral constructs. Actin was used as a loading control. ns, not significant; * *p* < 0.05; ** *p* < 0.01; *** *p* < 0.001

**Supplemental Table 1**: Detailed genotypes of all newly derived ES cell lines. The alleles used are LSL-Braf^V600E^ (Perna et al. 2015), LSL-Nras^Q61R^ (Burd et al. 2014), Pten^Flox^ (Lesche et al. 2002), Cdkna2^Flox^ (Krimpenfort et al. 2001), CAGs-LSL-rtTA3 (Dow et al. 2014), Tyr-CreERt2 (Bosenberg et al. 2006), and CHC (Beard et al. 2006).

**Supplemental Table 2**: List of primers used in this study.

Beard C, Hochedlinger K, Plath K, Wutz A, Jaenisch R. 2006. Efficient method to generate single-copy transgenic mice by site-specific integration in embryonic stem cells. *Genesis* **44**: 23–28.

Bosenberg M, Muthusamy V, Curley DP, Wang Z, Hobbs C, Nelson B, Nogueira C, Horner JW, Depinho R, Chin L. 2006. Characterization of melanocyte-specific inducible Cre recombinase transgenic mice. *Genesis* **44**: 262–267.

Burd CE, Liu W, Huynh MV, Waqas MA, Gillahan JE, Clark KS, Fu K, Martin BL, Jeck WR, Souroullas GP, et al. 2014. Mutation-specific RAS oncogenicity explains NRAS codon 61 selection in melanoma. *Cancer Discov* **4**: 1418–1429.

Dow LE, Fisher J, O'Rourke KP, Muley A, Kastenhuber ER, Livshits G, Tschaharganeh DF, Socci ND, Lowe SW. 2015. Inducible in vivo genome editing with CRISPR-Cas9. *Nat Biotechnol* **33**: 390–394.

Dow LE, Nasr Z, Saborowski M, Ebbesen SH, Manchado E, Tasdemir N, Lee T, Pelletier J, Lowe SW. 2014. Conditional reverse tet-transactivator mouse strains for the efficient induction of TRE-regulated transgenes in mice. ed. J.P. Lydon. *PLoS ONE* **9**: e95236.

Krimpenfort P, Quon KC, Mooi WJ, Loonstra A, Berns A. 2001. Loss of p16Ink4a confers susceptibility to metastatic melanoma in mice. *Nature* **413**: 83–86.

Lesche R, Groszer M, Gao J, Wang Y, Messing A, Sun H, Liu X, Wu H. 2002. Cre/loxP-mediated inactivation of the murine Pten tumor suppressor gene. *Genesis* **32**: 148–149.

Perna D, Karreth FA, Rust AG, Perez-Mancera PA, Rashid M, Iorio F, Alifrangis C, Arends MJ, Bosenberg MW, Bollag G, et al. 2015. BRAF inhibitor resistance mediated by the AKT pathway in an oncogenic BRAF mouse melanoma model. *Proc Natl Acad Sci USA* **112**: E536–45.

Xue W, Chen S, Yin H, Tammela T, Papagiannakopoulos T, Joshi NS, Cai W, Yang G, Bronson R, Crowley DG, et al. 2014. CRISPR-mediated direct mutation of cancer genes in the mouse liver. *Nature* **514**: 380–384.
