## Supplemental Data 1 for "A versatile ES cell-based melanoma mouse modeling platform"

Genotyping primers

***LSL-Braf V600E***
BRAFgeno F: ACTTTCATAGTCTAAGGGAGTTTGT
BRAFgeno R: CAAGATTCCTCTCCCAGCCG
WT: 300 bp, Mut: 230 bp; Primers Ratio 1:1

***LSL-NrasQ61R***
Nras-1: AGACGCGGAGACTTGGCGAGC
Nras-2: GCTGGATCGTCAAGGCGCTTTTCC
GENO2: GCAAGAGGCCCGGCAGTACCTA
WT: 487bp TG: 371 bp: Primer Ratio 1:1:1

***PTEN Flox***
PTEN genoF: CAAGCACTCTGCGAACTGAG
PTEN genoR: AAGTTTTTGAAGGCAAGATGC
WT: 160 bp Mut: 340 bp; Primer Ratio 1:1

***CDKN2A Flox***
CDKN2a genoFnew: CCTGACTATGGTAGTAAAGTGG
CDKN2a genoRnew: ACGTGTATGCCACCCTGACC
WT: 290 bp TG: 360 bp; Primer Ratio 1:1

***CHC***
C2 new 1: AATCATCCCAGGTGCACAGCATTGCGG
C2 new 2: CTTTGAGGGCTCATGAACCTCCCAGG
C2 new 6: GACGGCAATTTCGATGATG
WT: 250bp TG: 400bp; Primer Ratio 1:1:1

***Cags-LSL-rtTA3 (rosa26 kock-in)***
Rosa A: AAAGTCGCTCTGAGTTGTTAT
Rosa B: GCGAAGAGTTTGTCCTCAACC
Rosa C: CCTCCAATTTTACACCTGTTC
WT: 297bp TG: 350bp; Primer Ratio 2:1:2

***Tyrosinase-CreERt2***
CRE26: CCTGGAAAATGCTTCTGTGCG
CRE36: CAGGGTGTTATAAGCAATCCC
GABRA12: CAATGGTAGGCTCACTCTGGGAGATGATA
GABRA70: AACACACACTGGCAGGACTGGCTAGG
Cre: 400bp internal control: 300bp; Ratio 1:1:1:1

***CHC Targeted ESC***
PGK F: GAGCAGCTGAAGCTTATGGA
Hygro R: CTGAATTCCCCAATGTCAAG
oIMR7778: TATACTCAGAGCCGGCCT

oIMR7777: ACAGCGTGGTGGTACCTTAT
Targeted: 300bp internal control: 450bp; Primer Ratio 1:1:1:1

***Recombination of Braf exon 3 and 14 in ESC***
X3 F: CATGGCTTGAGTAAGTCTGC
X3 R: GATTCACATGGGACCTGAAC
X14 F: CTACCTAGTGAGACCATATCTC
X14 R: CAACAGTTGGATCCGTTTAAACG
WT: 400bp+250bp Recombined: 150bp; Primer Ratio 1.5:1

<u>T7 assay primers</u>

Pten Forward: CAGTCTCTGCAACCATCCAG
Pten Reverse: ATCTCATCTACACCCTAGAT

<u>SYBR Green primers</u>

rtTA3
Forward: TGACGACAAGGAAACTCGCT
Reverse: GGGGCAGAAGTGGGTATGAT
Product 134 bp

Mouse β-actin
Forward: TTGCTGACAGGATGCAGAAG
Reverse: ACATCTGCTGGAAGGTGGAC
Product 102 bp
